# Advancing scaffold biomimicry: engineering mechanics in microfiber scaffolds with independently controlled architecture using melt electrowriting

**DOI:** 10.1101/2023.05.28.542676

**Authors:** Brenna L. Devlin, Edmund Pickering, Mark C. Allenby, Naomi C. Paxton, Maria A. Woodruff

## Abstract

Melt electrowriting (MEW) is an additive manufacturing technique characterized by its ability to fabricate micronscale fibers from molten polymers into highly controlled 3D microfiber scaffolds. This emerging technique is gaining traction in tissue engineering and biofabrication research, however limitations in the ability to develop advanced coding to program MEW printers to fabricate scaffolds with complex fiber architectures has inhibited the development of structures with tunable and biomimetic mechanical properties. This study reports a series of non-straight scaffold architectures with combinations of independently controlled X & Y fiber spacing, corrections for MEW *jet lag*, and characterizations of their influences on scaffold mechanics. Polycaprolactone scaffolds with an elastic modulus ranging from 0.3 to 7.3 MPa were fabricated utilizing scaffolds manufactured from 5 layers of 55 μm fibers. The inclusion of scaffold design corrections in the gcode to compensate for decreasing deposition accuracy with increasing layer height enabled us to correct for discontinuous stress-strain mechanics and improved scaffold fabrication reproducibility. This study provides a comparison between a series of highly reproducible MEW scaffold architectures with non-straight fibers compared to the common crosshatch design to inform the development of more biomimetic scaffolds applicable to a variety of clinical applications. It further illustrates the significant effect toolpath correction has on reducing poor stress-strain mechanics, therefore improving the control, reproducibility, and biomimetic capacity of the MEW technique.

## 1. Introduction

Melt electrowriting (MEW) is revolutionizing biomaterial scaffold fabrication due to its ability to print highly ordered structures with small fiber diameters otherwise impossible to achieve with mainstream extrusion 3D printing techniques. As first described in 2011 [1], MEW uses pressure-driven extrusion coupled with high voltage to draw out microfibers of polymer melt from a nozzle onto a moving collector plate with high precision, shown in **Figure 1A**. In addition to allowing the fabrication of multi-scaled scaffold architectures and structures at high resolution, MEW is also gaining popularity for scaffolding applications in tissue engineering and regenerative medicine to support the growth of cells and tissue. These biomaterial scaffolds are key for mimicking the architecture of the extracellular matrix (ECM), the natural environment that provides structural and biomechanical support for cells. Most studies have thus far focused on simple box-structure meshes [2]–[6], herein referred to as *crosshatch*, due to the inherent simplicity, reproducibility and ease of fabrication. While often a necessary starting point, crosshatch scaffolds do not mimic the complex structures found in soft tissues and are unable to provide a realistic representation of composite tissue biomechanics.

**Figure 1.**
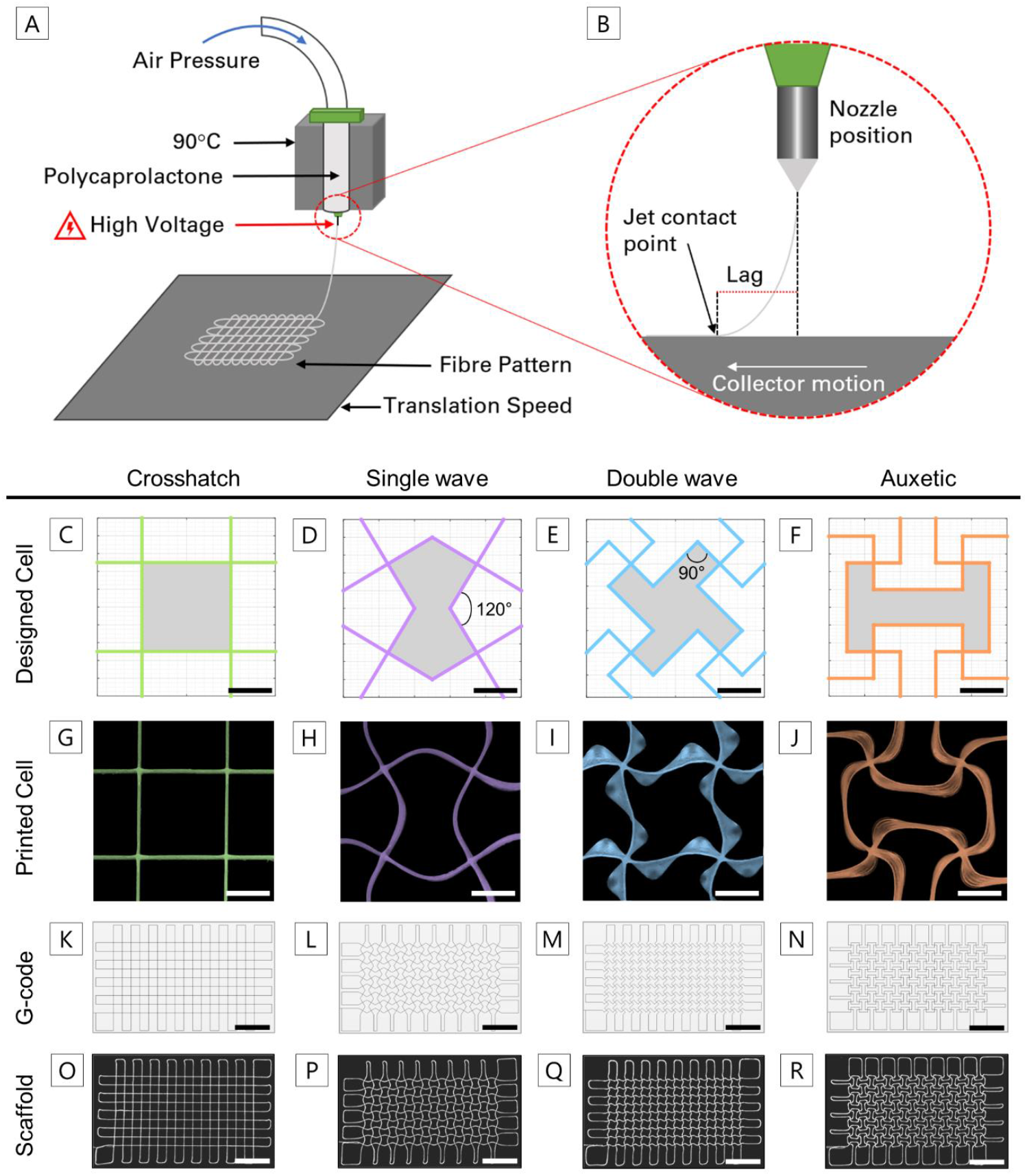
(A) Schematic diagram of the principles behind MEW and (B) a representation of the MEW *jet lag* effect experienced during printing; (C)-(F) schematic depiction of each geometry with the node junction of interest indicated by the red box and pore area shaded; (G)-(J) a snapshot of printed repeating cells represented with false colour; (K)-(N) the gcode visualization of each construct with (O)-(R) the corresponding printed construct. For (C-J) and (K-R), the scale is 750 μm and 6 mm, respectively.

In addition, MEW fabrication requires a single continuous fiber to create scaffold constructs which creates significant challenges in software programming and impacts the resultant mechanical properties. Better MEW scaffold design and printing software, known as gcode generation tools, are needed to determine which non-overlapping, non-straight fiber extrusion paths can be taken to manufacture more complex, biomimetic scaffold architectures [7] and to minimize printing artefacts such as those created by the discrepancy between the toolpath position and polymer contact point on the collector, herein referred to as MEW *jet lag*. MEW *jet lag* causes a mismatch between the nozzle position and jet contact point on the collector [8], [9], shown in **Figure 1B**. This is further complicated by material properties and the wide range of MEW operating parameters including collector plate translation speed, nozzle height, applied voltage and pressure. These software design and control improvements are necessary to unlock MEW’s full catalogue of clinical and industrial translational products. The rapid design and fabrication of biomaterial scaffolds that are representative of native tissues is an exciting area of research and is critical to develop efficient, scalable techniques for the clinical translation of tissue engineered constructs.

To this end, this study explored crosshatch scaffolds as the gold-standard in reproducible MEW fabrication and designed three scaffold geometries with non-straight fibers to create novel scaffolds with varying mechanical properties. Functionally, we present scaffolds with a *single wave* and *double wave* fiber geometry replacing the straight fibers in the 90 degree-intersecting crosshatch design as well as an *auxetic* scaffold design with significant elastic potential. These designs bear similarities to non-straight MEW scaffold designs presented in the literature in an effort to mimic specific tissue mechanics, including tendons and ligaments [10], [11] and cardiac tissue [12], [13], however these studies have extensively optimized subsets of MEW scaffold geometries for specific clinical requirements. There is a lack of analysis in the literature of how these geometries can be augmented and transformed to optimize scaffold mechanical behaviour. Specifically, the role of independently controlling X & Y spacing of such straight and non-straight fiber scaffolds has not yet been explored, nor the impact of correcting for fabrication artefacts such as layer stacking anomalies on scaffolds with curved geometry. This study therefore seeks to provide a systematic investigation of MEW scaffold geometries to inform the development of biomaterial scaffolds that mimic the mechanics of the native tissues they seek to augment or replace, consolidating observations of scaffold printing artefacts [13], [14] with ‘smart’ scaffold design capability to produce for the first time a systematic exploration into the relationship between scaffold design and mechanics.

## 2. Results and Discussion

### 2.1. Scaffold Design & Fabrication

To assess the effects of introducing non-straight fibers on scaffold mechanics, scaffold structures were designed in MATLAB and fabricated using the MEW process as described previously [14] and shown in **Figure 1A**. Four scaffold structures were investigated, shown in **Figure 1C-F**. These are referred to as *crosshatch*, which featured overlapping orthogonal layers of straight fibers **(Figure 1C)**, and three non-straight designs; a *single wave* with a wave angle of 120° **(Figure 1D)**, a *double wave* with a wave angle of 90° **(Figure 1E)** and an *auxetic* which utilizes a ‘dog-bone’ shape to promote a negative Poisson’s ratio **(Figure 1F)**. 5-layer 24 × 12 mm rectangular scaffolds were successfully and reproducibly fabricated with an average fiber diameter of 54.5 ± 0.6 μm (N = 4 scaffold designs, n = 3 scaffold replicates). The inclusion of turn-around regions [15] outside the margins of the programmed scaffold **(Figure 1K-N)** were not included in the scaffold dimensions but were necessary to allow the continuous fiber to move to the next section without affecting the resulting construct. **Figure 1G-J** shows a comparable section of the scaffold to the programmed print design and demonstrates the varying levels of fiber deposition accuracy achieved when introducing non-straight fibers. In MEW, *jet lag* error is expected [16] and causes a deviation between the programmed and true fiber laydown path as depicted in **Figure 1B**. This was observed through the loss of discrete vertices which resulted in smoother turns rather than sharp changes in direction (shown in **Figure 1O-R)**. Note, later in this work, we explore the impact of MEW *jet lag* error on scaffold mechanics and explore correction methodologies.

### 2.2. Influence of Patterning on Scaffold Mechanics

The scaffold geometry plays a key role in determining the mechanical properties of the resulting construct. For instance, biological tissues often exhibit a J-shaped stress-strain response with three distinct phases [13], [17]; (1) a low-stiffness non-straight elastic region (toe) dominated by fibers aligning to the loading direction, followed by a transition (heel) into (2) a linear higher-stiffness region, before (3) eventual plastic deformation. MEW unlocks the potential for near unlimited variation in mechanics by varying the print pattern. This can be exploited to create MEW scaffolds that mimic these natural tissue mechanics. To characterize the mechanical behaviour, scaffolds were tensile tested in the longitudinal direction (loading direction) as shown in **Figure 2A**. The stress-strain curves for each group are shown in **Figure 2B**.

**Figure 2.**
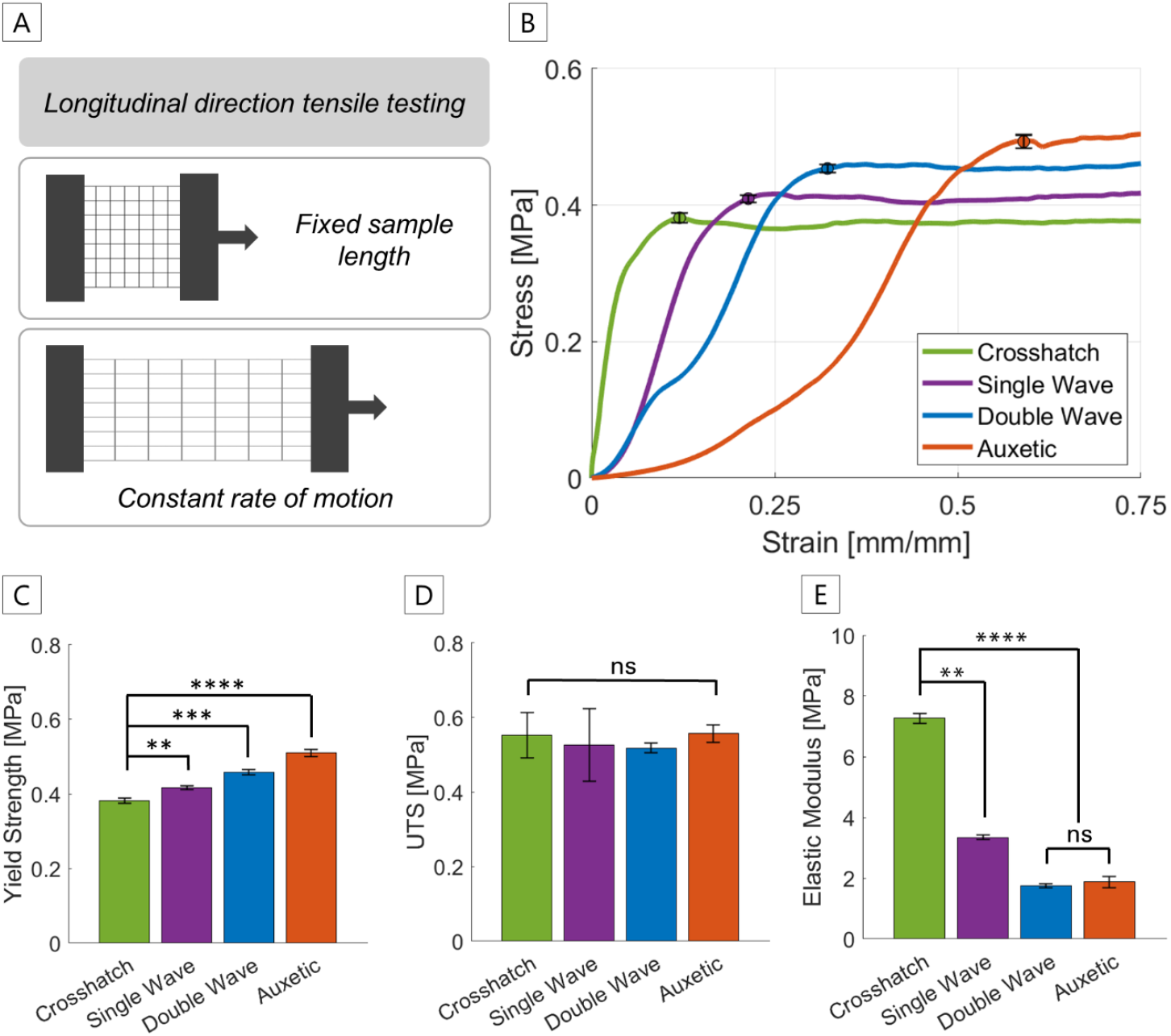
(A) A schematic representation of the tensile testing procedure (B) the stress-strain curves for each of the scaffold geometries and illustrating their standard deviation around one representative stress-strain point (n = 3). Comparison of key mechanical properties for each unit cell tested; (C) yield strength, (D) ultimate tensile strength and (E) elastic modulus. **p<0.001, ***p < 0.0001, ****p < 0.00001 and ns = non-significant.

The *crosshatch* scaffold, comprised of straight fibers already aligned to the loading direction, demonstrated high stiffness with no initial toe region. The *crosshatch* did not exhibit a J-shape response, which makes it a poor design for replicating tissue mechanics. The fabrication of non-straight fibers for the *single wave, double wave* and *auxetic* structures saw the introduction of features associated with the J-shaped stress-strain curve reminiscent of biological soft tissues, including an exaggerated toe and heel prior to the linear elastic region. These toe and heel regions are more prominent in the geometries featuring longer lengths of crimped fibers, and therefore longer strain range over which it uncrimps.

To further illustrate differences in mechanics, the yield strength, ultimate tensile strength (UTS) and elastic modulus were calculated. The addition of longer lengths of crimped fibers increased the elastic region prior to yielding (**Figure 2C**), with the constructs able to withstand a higher strain prior to permanent deformation. No significant difference was observed in the UTS (**Figure 2D**) between the groups. This is explained by all the longitudinal fibers having aligned to the loading direction with deformation being governed by axial elongation of the fibers, similarly in all scaffolds. A significant reduction in elastic modulus (**Figure 2E**) corresponding to the increased toe and heel regions was observed with an approximate 55-76% reduction between the *crosshatch* and non-straight designs. This illustrates an ability to manipulate and target specific construct mechanics from the conceptualisation phase prior to fabrication. In addition, the *double-wave* and *auxetic* scaffolds demonstrated a shifting between successive layers (see **Figure 1I-J**) where successive layers are not perfectly stacked, resulting from MEW *jet lag*. This has a significant effect on the mechanics, causing two linear regions (in the case of the *double wave*) or causing an exaggerated heel region (for the *auxetic*). The cause and effect of this is explored further in Section 2.4.

### 2.3. Influence of Geometric Non-Uniformity on Scaffold Mechanics

The next objective of this work was to explore the effects of scaffold geometry on mechanics through independent control of X & Y fiber spacing in the longitudinal or transverse direction. In addition, control of the fiber spacing essentially ‘stretched’ the geometry in the relevant direction, affecting the wave angles and creating constructs with anisotropic mechanics. This design consideration has the potential to create numerous constructs from a single pattern, however in this work we present two non-uniform variations of the original geometry as shown in **Figure 3**, and subsequently defined as cases A, B & C. This resulted in *original constructs with no change (Case A), half the number of fibers in the transverse direction (Case B)*, or *half the number of fibers normal to the direction of loading (Case C)* to assess the difference in mechanical properties with an irregular deformation of the programmed pattern. Tensile testing of the non-straight groups revealed several interesting patterns when compared back to the *original constructs*.

**Figure 3.**
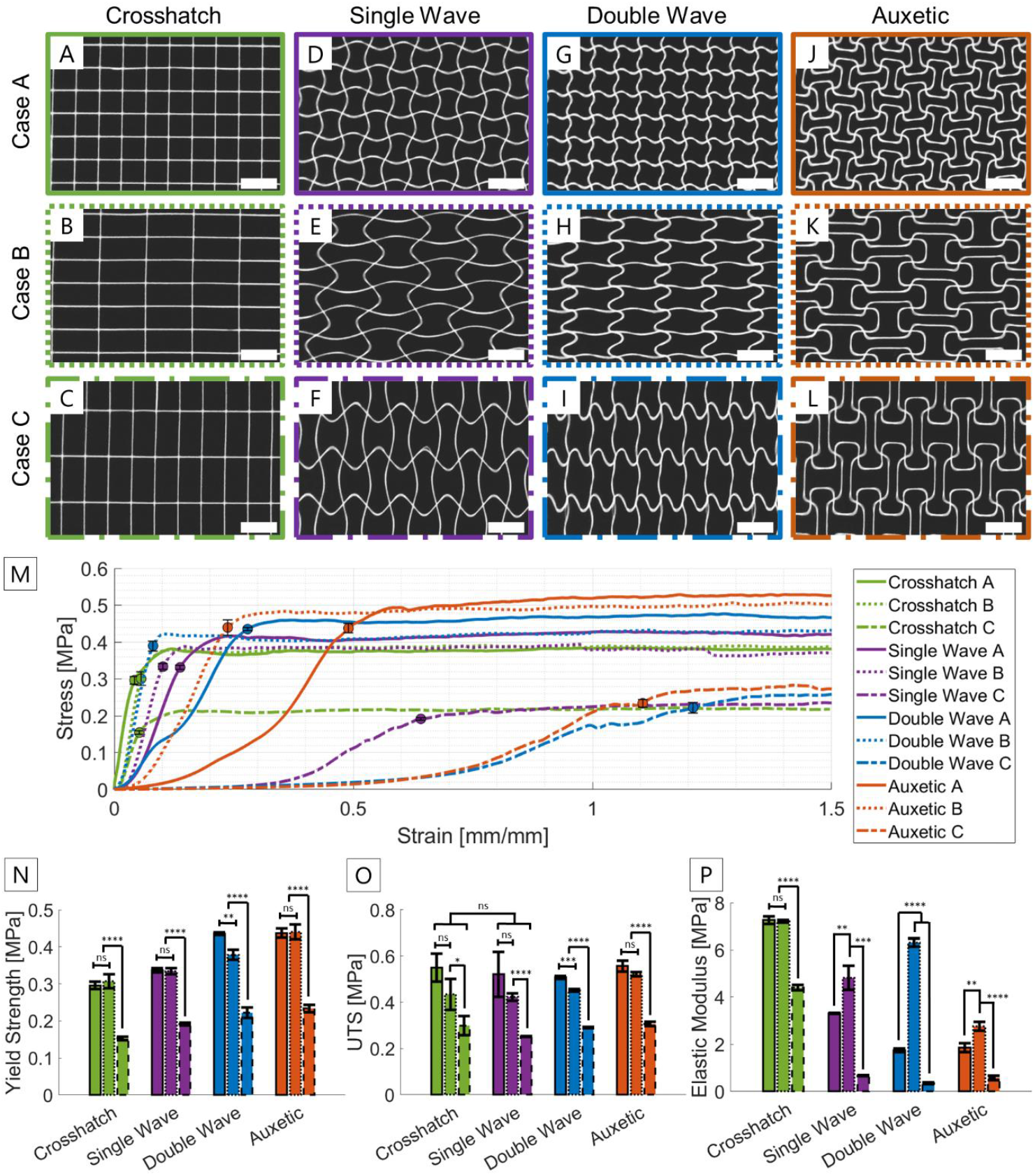
Representative images of scaffold constructs in each print configuration; no change (case A), deformed in the longitudinal direction (case B) and transverse direction (case C); (A)-(C) Crosshatch; (D)-(F) single wave; (G)-(I) double wave; and (J)-(L) auxetic geometry. A summary of key mechanical properties for each unit cell configuration tested; (N) yield strength, (O) ultimate tensile strength and (P) elastic modulus. Scale is 5 mm. *p < 0.01, **p < 0.001, ***p < 0.0001, ****p < 0.00001 and ns = non-significant.

*Half-transverse constructs (Case B)*, as depicted in **Figure 3B, E, H & K** showed no significant change in the *crosshatch* elastic modulus, due to the number of loaded fibers (longitudinal fibers) remaining unchanged. Conversely, the stiffness of all other designs compared to *original constructs (Case A)* was shown to increase significantly. The loaded fibers in this case were elongated and featured less curvature, resulting in a reduction of MEW *jet lag* error. No significant change in UTS and yield strength was observed for the *crosshatch, single wave* and *auxetic* designs, however the *double wave* design did experience a small but significant change in these properties resulting from MEW *jet lag* inconsistencies. The non-straight constructs do not lose their mechanical biomimicry of soft biological tissues and maintain the J-shape stress-strain curve and overall, the *half-transverse case* demonstrates an ability to manipulate construct elasticity and pore area without significantly impacting mechanical yield and UTS.

*Half-normal constructs (Case C)* as shown in **Figure 3C, F, I & L** caused an expected drop of 46% in UTS and corresponding reduction of 56% in yield strength compared to *original constructs*, due to the halved number of fibers in the loading direction. A significant reduction of elastic modulus was observed, with non-straight groups showing an increased low stiffness non-straight toe and heel region when compared to *original constructs*. In all designs, the slower transition to the linear elastic region caused a significant reduction in elastic modulus and the reduction of fibers caused an expected drop in yield strength. This is significant as this behaviour increased the strain which the sample was able to withstand before permanent deformation, and the predictable changes in UTS and yield strength have implications for reducing the iterative nature of scaffold design and fabrication. The stress-strain data for each group is shown in **Figure 3M**, with the average yield strength, UTS and elastic modulus represented in **Figure 3N, O & P**. Utilizing the capability of MEW to create non-straight constructs is gaining popularity in the literature [11], [12], [18] and characterizing the mechanical potential of such constructs is paramount to furthering MEW’s promise as a tissue engineering technology.

### 2.4. Effects of Tool Path Correction on Scaffold Mechanics

One of the unfortunate hallmarks of MEW is a jet lag effect, caused by a mismatch between the printing nozzle and polymer contact point on the collector and resulting in reduced fiber deposition accuracy and stacking errors which become more pronounced with multiple layers or the introduction of non-straight scaffold architecture [9], [16], [19]–[21]. Not only does this impact the structure, but this also has significant impacts on the mechanics, as touched on briefly in the previous sections. It thus becomes relevant to develop techniques to correct this. To counter this MEW *jet lag* effect and correct for stacking errors, the midpoint coordinate of each curve was systematically offset by 4-5% for each layer, similar to existing methods reported in the literature [9], [16], [22]. The scaffolds were then imaged and analyzed in ImageJ to quantify the difference in fiber length between first and last print layers.

In the case of the *double-wave* scaffold, the effects of MEW *jet lag* (**Figure 4A**) resulted in top-layer fibers being 25% shorter than the bottom layer and only approximately 20° away from being completely straight (see **Supplementary Information, section A**). The programmed tool path offset which resulted in a significant improvement in maintaining the geometric morphology through corrected fiber placement can be observed in **Figure 4B**. The effect of this is that fibers in the upper layer both straighten and yield earlier than the longer fibers in the lower layers. In other terms, fiber yielding does not occur at the same strain in each layer, rather these occur sequentially (the same occurs for the heel). This creates the bi-linear deformation behavior seen in **Figure 4C**. This behavior of continually shortening fibers is expected to become more significant for taller scaffolds as the fibers gradually tend towards becoming straight.

**Figure 4.**
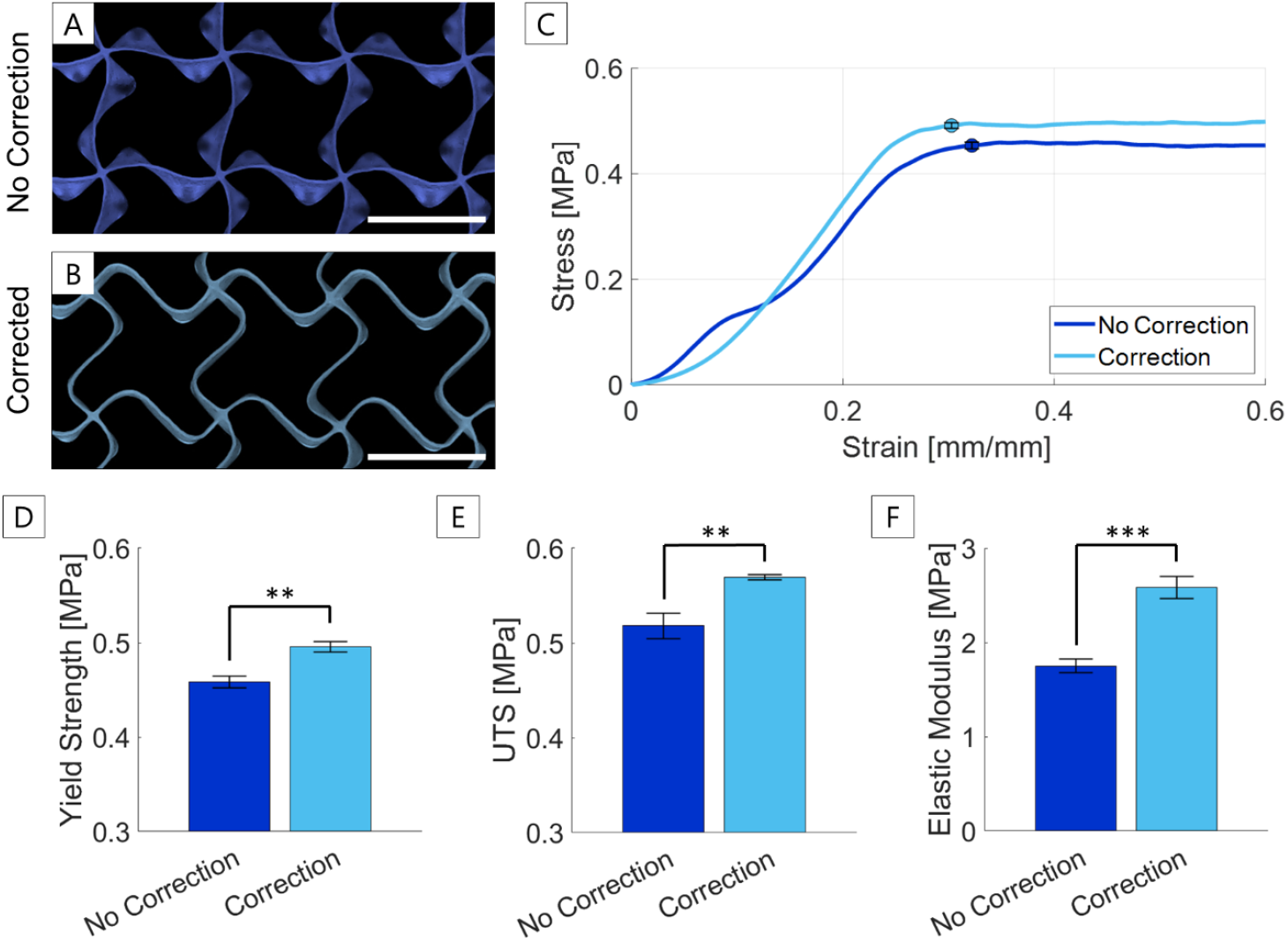
Effect of toolpath correction on microfiber scaffold mechanics. Printed construct represented in false colour with no toolpath correction (A), lag correction (B) and the linear elastic region of the mechanical response data (C). A comparison of key mechanical properties between the uncorrected and corrected constructs, the 0.2% offset yield (D), ultimate tensile strength (E) and elastic modulus (F). Scale is 5 mm; **p<0.001, ***p < 0.0001.

Following correction, minimum variation is observed between individual layers. This manifests in the mechanics, by a significantly improved J-shape stress-strain behavior. While in the uncorrected case a bi-linear behaviour was observed due to the differing fiber length, in the corrected case each layer fiber is equal length, resulting in a J-shape stress-strain behavior, like that of native tissue. The yield point was shown to significantly increase (**Figure 4D**) with equal lengths of fibers. Correcting for MEW *jet lag* does not just provide a smoother stress-strain curve, but also improves the mechanics, increasing elastic modulus and preventing premature yield, with the corrected scaffold having a 37% increase in elastic modulus (**Figure 4E**) and 28% increase in ultimate tensile strength (**Figure 4F**), respectively. A modest correction of 4-5% per layer was sufficient to eliminate the observed bi-linear elastic region and vastly improve the mechanical properties, demonstrating the significance of gcode toolpath correction to account for fiber placement inaccuracies resulting from MEW *jet lag*.

## 3. Conclusion

In this study, we conducted a comprehensive analysis of four different scaffold architectures fabricated using MEW. The X and Y fiber spacing in each scaffold design were independently controlled, allowing us to manipulate their mechanical properties based on straightforward geometric considerations. Additionally, we investigated the impact of MEW polymer fiber correction on the overall mechanics of the scaffolds.

The non-straight scaffolds presented in this study exhibit a wide range of mechanical properties, and unlike the typical *crosshatch* pattern, showcase distinct J-shaped stress-strain curves that resemble those observed in biological tissues. The MEW scaffold architecture and the introduction of non-uniform deformation in the programmed pattern were found to greatly influence scaffold mechanics. In particular, all non-straight groups exhibited enhanced elasticity and an expanded strain region, surpassing the performance of the *crosshatch* pattern. MEW *jet lag* had a considerable negative impact on non-straight fiber placement, with top-layer fibers being 25% shorter than the bottom layer, causing uneven fiber yielding and a bi-linear stress-strain response. To overcome inaccuracies in fiber placement, we implemented a straightforward percentage offset approach, resulting in improved and more consistent scaffolds. These corrections can have profound implications for the mechanical characteristics and suitability of scaffolds for their intended purpose; for example, bone implants need high stiffness to bear weight during the amelioration period [23] which is very different to those for ligament repair which require more elasticity and the ability to recover cyclically from high straight mechanobiological loading [24]. As the ‘gold standard’ geometry in reproducible MEW scaffold fabrication, the *crosshatch* becomes less desirable due to its simplicity and inability to represent native tissue structures.

More complex scaffolds designs will inevitably emerge as the MEW technology improves. This study demonstrates the ability to manipulate the scaffold structures of multiple non-straight MEW designs for targeted manipulation of mechanical properties. Moreover, advancements in gcode generation software and MEW-specific tool path corrections facilitate the rapid conceptualization and generation of accurate and reproducible tool path patterns. These design manipulations, facilitated by a custom MATLAB program, offer a new level of design freedom for the efficient production of biomimetic MEW scaffolds. The ability to print highly ordered structures with micron-scale precision cements MEW technologies as a promising avenue for fabricating biomimetic scaffolds for use in wider tissue engineering applications.

## 4. Methods

### Scaffold Design & Fabrication

Scaffold cells were designed in MATLAB using a custom gcode generation library. 5-layer, 24 × 12 mm rectangular scaffolds were fabricated using four scaffold structures, referred to *crosshatch, single wave, double wave* and *auxetic* (shown in Figure 1C-F). Turn-around regions outside the margins of the programmed scaffold (Figure 1K-N) were not included in the scaffold dimensions. Polycaprolactone (45 kDa, Sigma Aldrich) was heated to 90°C and printed using a custom-built MEW printer described previously [14], [25] using a 21 G needle. A working distance of 4.5 mm, air pressure of 0.05 MPa and a translation speed of 600 mm/min was maintained for each print. Voltage was adjusted by up to 0.4 kV to minimize variation in fiber diameter. Scaffolds were printed at 22.5°C ambient temperature and 48% relative humidity.

### MEW Toolpath Correction

A layer-by-layer offset for the printing path amplitude described by Liashenko et al. (2020) [16] was used to correct for the MEW *jet lag* which causes a tendency for nonlinear fibers to straighten in each consecutive layer. The percentage offset for each design was determined through trial and error until a visually corrected scaffold was achieved and for all cases was found between 4-5%.

### Imaging & Analysis

The diameter of individual fibers (12 total, N = 4 scaffold groups, n = 3 scaffold replicates) was captured using an optical microscope (Axio Observer 7, Zeiss) with a digital collector using a 5X objective. Imaging of complete and magnified scaffolds was captured using an iPhone 11 digital camera and a Digicom digital microscope, respectively. ImageJ was used to analyze fiber lengths and laydown angles between first and final layers of printed *double wave* constructs to investigate the effects of MEW toolpath correction on fiber deposition. Fiber length was estimated by fitting a spline between fiber junctions (n = 12) and fiber laydown angle by the approximate angle between the back and forward tangents of the curve (n = 14).

### Mechanical Testing

Samples in triplicate from each scaffold design underwent mechanical testing (Tytron 250, MTS Systems) in triplicate with a 10 N load cell under displacement control. Three separate experiments were performed: (A) A comparative analysis of each design (*crosshatch, single wave, double wave* and *auxetic*) using a total of 12 samples (N = 4, n = 3), (B) Two perturbations of the original design (N = 2, n = 3); half the number of fibers in the transverse direction or normal to the direction of loading, referred to as the *half-transverse (Case B)* and *half-normal (Case C)*, respectively, and (C) an analysis of MEW toolpath correction on the resulting mechanics of the *double wave* design (N = 2, n = 3).

Stress was calculated as the tensile force over cross-sectional area with a width of 12 mm as defined by the print pattern and an average scaffold height determined through the fiber diameter multiplied by the total number of layers. The strain was determined based on a scaffold length of 10 mm as defined by the sample test length and ultimate tensile strength via the maximum stress in the stress-strain curve. Elastic modulus was determined via the gradient of the linear region, following the heel. In the *double wave* case, which showed two defined linear regions, the elastic modulus and yield stress was estimated over the total region. The yield point was approximated as the point in the stress-strain curve at which the curve levels off and exhibits a zero slope as defined in ASTM D638.

### Statistics

All data is reported as mean ± standard deviation from replicate samples. Statistical analysis was conducted in MATLAB. Student’s t-test and one-way ANOVA were used to determine significance between two and three groups, respectively. In all cases, statistical significance was determined by a *p* value of less than 0.05.

## Supporting Information

Supporting Information is available from the Wiley Online Library or from the author.

## Acknowledgements

The authors acknowledge the continued support from the Queensland University of Technology. These experiments were conducted using the resources of the Central Analytical Research Facility within the Research Infrastructure Division, Queensland University of Technology. B.L.D is supported by a Queensland University of Technology Postgraduate Research Award. N.C.P acknowledges support from an Advance Queensland Industry Research Fellowship (AQIRF), QUT Early Career Research Scheme grant and the Knight Campus-PeaceHealth Postdoctoral Fellowship Program. M.C.A acknowledges support from the Australian Research Council (DECRA fellowship DE220100757). Edmund Pickering was supported via a QUT Centre for Biomedical Technologies research fellowship.

