## Supplementary Information 1 for "Advancing scaffold biomimicry: engineering mechanics in microfiber scaffolds with independently controlled architecture using melt electrowriting"

### Supporting Information: fiber length and angle measurements

A **Fiber length**

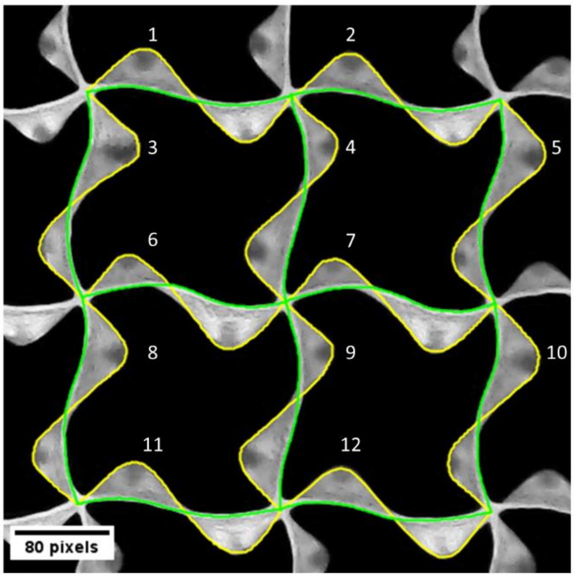


**Yellow:** first print layer
**Green:** final print layer

| Fiber length (pixels) | | |
|---|---|---|
| **#** | **First layer** | **Final layer** |
| **1** | 239.20 | 179.26 |
| **2** | 236.23 | 178.25 |
| **3** | 234.91 | 176.42 |
| **4** | 234.67 | 179.50 |
| **5** | 238.42 | 177.03 |
| **6** | 237.88 | 174.48 |
| **7** | 237.27 | 180.35 |
| **8** | 238.35 | 177.59 |
| **9** | 245.74 | 179.78 |
| **10** | 237.39 | 179.75 |
| **11** | 237.20 | 177.35 |
| **12** | 233.61 | 182.80 |
| **Mean** | **237.55** | **178.53** |
| **St Dev** | **3.08** | **2.17** |

B **Fiber angle**

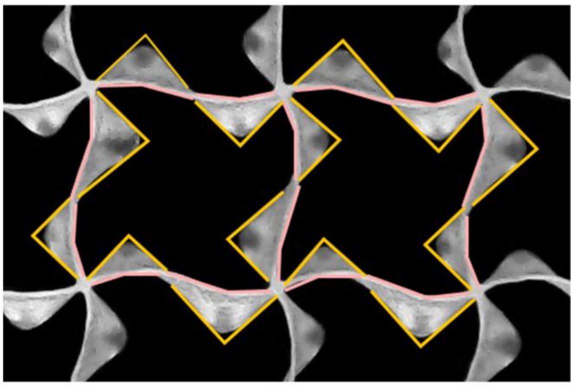

**Yellow:** first print layer
**Pink:** final print layer

| Fiber angle (degrees) | | |
|---|---|---|
| **#** | **First layer** | **Final layer** |
| **1** | 85.56 | 161.67 |
| **2** | 88.93 | 160.95 |
| **3** | 86.34 | 160.56 |
| **4** | 86.10 | 160.21 |
| **5** | 82.85 | 157.77 |
| **6** | 89.10 | 153.61 |
| **7** | 90.84 | 159.15 |
| **8** | 91.34 | 155.50 |
| **9** | 89.53 | 156.11 |
| **10** | 86.50 | 159.05 |
| **11** | 87.41 | 165.58 |
| **12** | 88.55 | 155.65 |
| **13** | 89.00 | 161.91 |
| **14** | 89.91 | 162.98 |
| **Mean** | **87.97** | **159.30** |
| **St Dev** | **2.32** | **3.32** |

**Figure 1 –** Supporting information for Section 2.4: Effects of tool path correction on scaffold mechanics. (A) Fiber length (n = 12) was estimated by fitting a spline over the first (yellow) and final (green) layers of the *double wave* scaffold construct. Fiber length demonstrated a 25% reduction in length from first to final layer. (B) Fiber angle (n = 14) was estimated using the angle tool with laydown angles of the first and final layers indicated in yellow and pink, respectively. The laydown angle increased by approximately 81% from the first to the final layer, demonstrating a significant loss in accurate fiber placement. For (A-B), each number is a separate measurement of the superimposed lines.
